# A land plant specific VPS13 mediates polarized vesicle trafficking in germinating pollen

**DOI:** 10.1101/2022.11.01.514778

**Authors:** Surachat Tangpranomkorn, Motoko Igarashi, Fumiko Ishizuna, Yoshinobu Kato, Takamasa Suzuki, Sota Fujii, Seiji Takayama

## Abstract

Pollen has an extraordinary ability to convert from a dry state to an extremely rapidly growing state. During pollination, pollen receives water and Ca^2+^ from the contacting pistil, which will be a directional cue for pollen tube germination. The subsequent rapid activation of directional vesicular transport must support the pollen tube growth, but the molecular mechanism leading to this process is largely unknown. Here we show that a plant-specific VPS13, AtVPS13a, mediates vesicle trafficking during the polarization process in *Arabidopsis* pollen. *AtVPS13a* knockout severely affected pollen germination and lipid droplet discharge, while Ca^2+^-dynamics after pollination was unchanged. Cellular distribution patterns of AtVPS13a and a secretory vesicle marker were synchronized, with a slight delay to the Ca^2+^-dynamics in polarizing pollen. The absence of AtVPS13a led to reduced cell wall deposition during pollen germination. These results suggest that AtVPS13a mediates pollen polarization, by regulating proper directional vesicular transport following Ca^2+^-signaling.

## INTRODUCTION

Pollen usually produced as a quiescent unit but rapidly establish cell polarity upon pollination to germinate and deliver sperm cells towards an ovule. In plant species with dry stigmas, including the model plant *Arabidopsis thaliana*, pollen coat is mixed with stigma pellicle to form “foot” structure at the pollen-stigma contact. The structure functions as a conduit for the transfer of necessary liquid for pollen germination (Swanson et al., 2004). Pollen hydration reactivates cellular metabolism and induce cell polarity by possible cues such as directional water supply or local Ca^2+^ influx, which ultimately leads to pollen tube outgrowth at the stigma contact site (Wolters-Arts et al., 1998; Lush et al., 1998; Iwano et al., 2004; Chen et al., 2013).

Pollen cells experience drastic physiological changes before germination. Pollen uptake Ca^2+^ after hydration and form intracellular gradient adjacent to the pollination site, and pollen grains fail to germinate without this local Ca^2+^ elevation (Iwano et al., 2004). Asymmetric filamentous actin (F-actin) network also mediates pollen grain polarity. Common F-actin pattern found in germinating pollen of many species, including *Arabidopsis*, is the radially aligned F-actin towards pollen aperture or future germination site (Heslop-Harrison and Heslop-Harrison, 1992; Gibbon et al., 1999; Zhang et al., 2011; Liu et al., 2018). More specifically, this collar like F-actin structure is orchestrated by *Arabidopsis* Formin homology5 (AtFH5) which relocates onto plasma membrane at the polarized site and nucleates actin assembly (Cheung et al., 2010; Liu et al., 2018). The translocation is thought to bring small vesicles to the polarized site (Liu et al., 2018). In *Arabidopsis* pollen, secretory vesicles gather at the germination site at around 20 minutes after pollination (Kandasamy et al., 1994). Octameric exocyst complex mediates the exocytosis of vesicles and facilitates the trafficking of cell wall materials for the formation of germination plaque (Hoedemaekers et al., 2015; Li et al., 2017). Although various cellular events have been reported during pollen polarity establishment, the molecular mechanism leading to such process is still missing.

Several genetic screens involving visible phenotypes such as male sterility (Preuss et al., 1993; Hulskamp et al., 1995) or distorted segregation (Johnson et al., 2004; Boavida et al., 2009; Hoedemaekers et al., 2015) have been used to isolate mutants with defects in reproductive process, however, molecular mechanisms specific to the early stages after pollination have rarely been revealed. In this study, we established a forward genetic screening scheme to identify mutants defective in early events after pollination. We found that AtVPS13a mediates pollen polarization by regulating organelle rearrangements and ensuring sufficient vesicle trafficking for pollen germination. AtVPS13a showed polarized localization in germinating pollen grains with exocyst subunits or the secretory vesicle marker. We discuss the possibility that AtVPS13a is the early pollination signal transducer after Ca^2+^ influx, which establishes pollen polarity by guiding vesicle secretion to the pollen germination site.

## RESULTS

### Search for novel pollen factors involved in the compatible pollination process

Previously, we performed a transcriptomic analysis and identified 398 genes that are induced in stigmas after pollination with compatible pollen (Iwano et al., 2014). *Dicarboxylate carrier 2* gene (*DIC2*) was up-regulated 12.1-fold after pollination, and we fused its promoter with the *Photinus pyralis* luciferase gene. This reporter construct was introduced into the wild-type (WT) *A. thaliana* strain Col-0 to generate a system for using reporter pistils in a 96-well plate assay (Supplemental Figure S1; see Methods for details). WT pollen grains induced bioluminescence in this system, but those from the known pollen coat mutants *cer1* (Hulskamp et al., 1995), *cer6-2* (Preuss et al., 1993), and *fkp1* (Ishiguro et al., 2010) did not (Figure 1A). This indicated that this novel luciferase bioluminescence-based system can specifically detect pollen-side mutations involved in the early pollination process.

**Figure 1.**
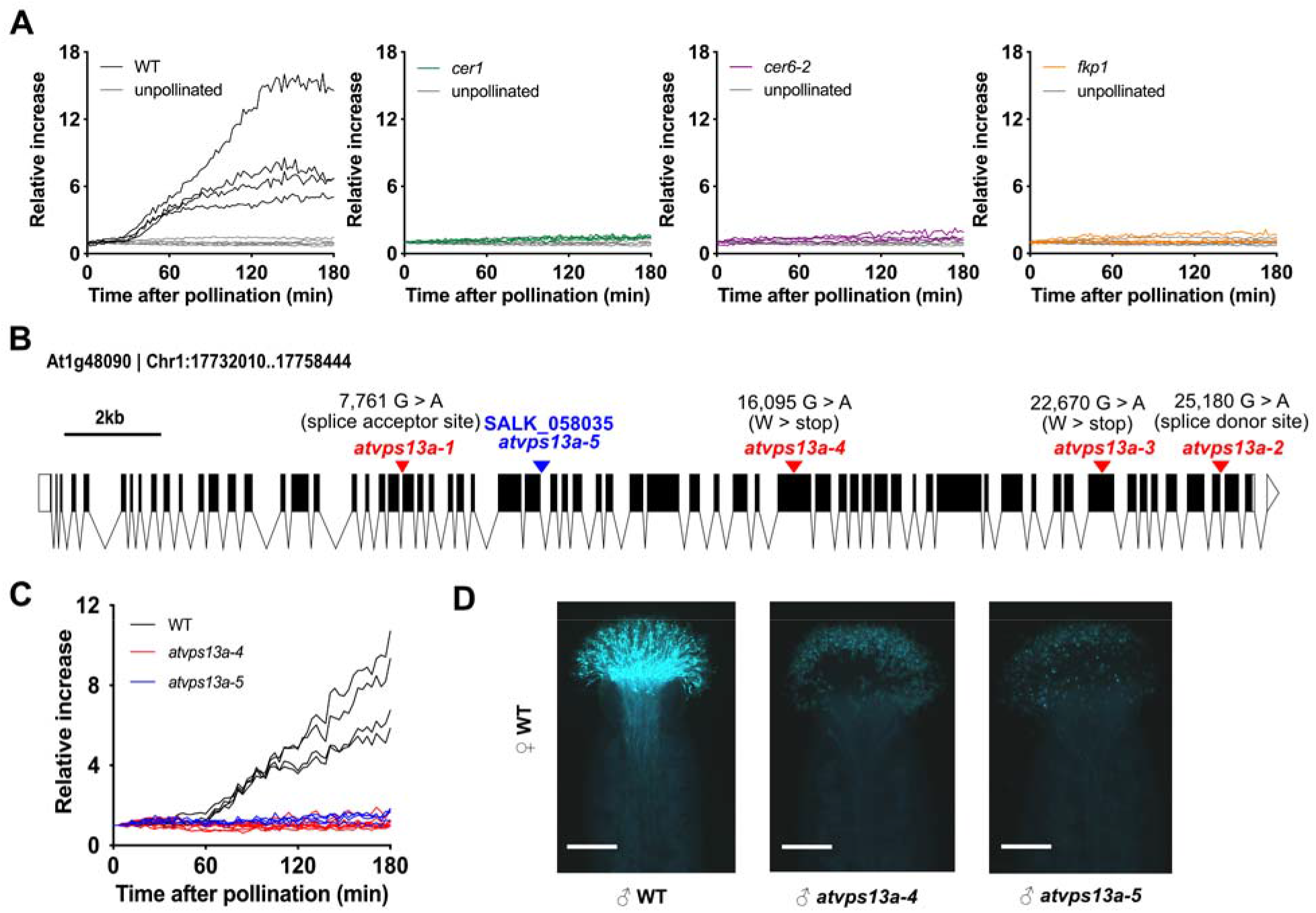
At1g48090 is the causal gene of the non-induction mutant phenotype. (**A**), Relative increases in the luciferase activity of reporter pistils after pollinated with WT and pollen coat mutants (*cer1, cer6-2* and *fkp1*) pollen, one line represents one reporter pistil. At least 3 pistils were pollinated by a pollen donor of each genotype. (**B**), Gene structure of At1g48090. The exon-intron backbone was generated on http://wormweb.org/exonintron. The arrowheads indicate the locations of the EMS-generated mutations (red) and the T-DNA insertion (blue). *atvps13a-1* has a nucleotide change at a 3’ splice acceptor site, *atvps13a-2* has a nucleotide change at a 5’ splice donor site, and *atvps13a-3* and *atvps13a-4* contain nonsense mutations. (**C**), Luciferase-based pollination assays with WT and *atvps13a* mutant pollen. (**D**), Aniline blue staining of pistils after pollination with WT or mutant pollen for 3 h, bars = 200 μm.

We screened the pollen of approximately 2,400 M_2_ individuals obtained from the ethyl methanesulfonate (EMS) mutagenesis and identified 11 independent pollen mutants. Four of them did not induce stigmatic bioluminescence and seven were reduced in induction compared to WT pollen. We focused our study on the non-induction mutants and used the bulked-segregant analysis with Mitsucal pipeline (Suzuki et al., 2018) to identify the causal mutations (see Methods for details). All the non-induction mutants carried either a nonsense mutation or a mutation in a splice acceptor/donor site within the gene At1g48090 (*atvps13a-1–4*; Figure 1B, Supplemental Figure S2). In agreement with these EMS-generated mutants, pollen from a T-DNA insertion mutant (SALK_058035, *atvps13a-5*) failed to induce bioluminescence in reporter pistils and did not germinate efficiently on stigmas (Figure 1C, D). These results indicated that the At1g48090 gene is responsible for the non-induction phenotype. For further experiments, we used *atvps13a-5*, referred to here as *atvps13a* unless stated otherwise.

### AtVPS13a is a land plant unique form of VPS13

The At1g48090 gene was predicted to encode a 4,132 amino acid calcium-dependent lipid-binding protein with similarity to the yeast Vacuolar Protein-Sorting associated protein13 (VPS13, GenBank: AJV75928.1). In addition, we also found two other *A. thaliana* genes encoding VPS13 (At4g17140 and At5g24740). A phylogenetic analysis of VPS13 from different eukaryotic organisms revealed that the At5g24740 protein, named SHRUBBY in a past study (Koizumi and Gallagher, 2013), may be the ancestral form (Figure 2A), since its molecular size resembles yeast and human VPS13A (Figure 2B). At1g48090 and At4g17140 (*AtVPS13a* and *AtVPS13b*, respectively) belong to a subgroup that specifically diverged in the land plant lineage (Figure 2A). All *Arabidopsis* VPS13s maintained VPS13 structural domains at N-terminus and C-terminus found in both human HsVPS13A and yeast ScVPS13, suggesting that VPS13 protein structure are broadly conserved (Figure 2B). VPS13 family members are generally thought to be involved in lipid transport between organelles (Leonzino et al., 2021). The hydrophobic cavity structure at the N-terminal amino acids involved in lipid transport function (Kumar et al., 2018). These N-terminal cavities with approximately 300 amino acids were conserved in the modeled structures of *Arabidopsis* VPS13s (Supplemental Figure S3A). An amphipathic helix near the C-terminus of HsVPS13A, which localizes to the lipid droplets (Kumar et al., 2018), was also likely conserved in *Arabidopsis* VPS13s (Supplemental Figure S3B). On the other hand, only AtVPS13a had a putative C2 domain, which generally is involved in Ca^2+^-dependent protein targeting to biological membrane (Corbalan-Garcia and Gómez-Fernández, 2014) (Figure 2B). The predicted protein structures suggest that AtVPS13a may have acquired specific domains that might be essential to support land plant-specific biological event.

**Figure 2.**
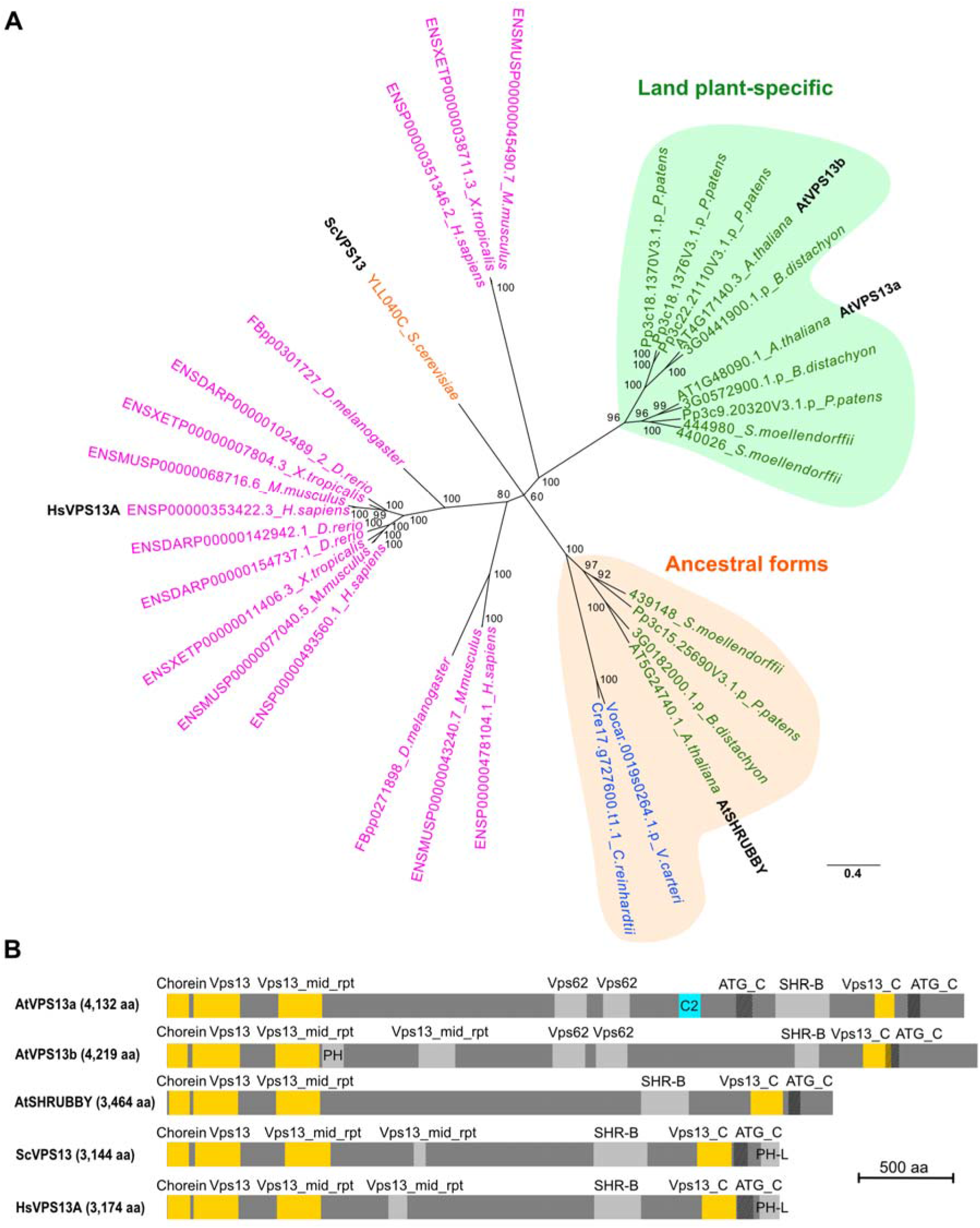
AtVPS13a and AtVPS13b are land plant-specific VPS13 isoforms with different domains in the middle parts of the proteins. (**A**), Unrooted phylogenetic tree of VPS13 proteins: magenta, animals; orange, yeast; green, land plants; blue, green algae. The scale bar indicates substitutions per site. (**B**), Gene structure of the *Arabidopsis* VPS13s in comparison with yeast ScVPS13 and human HsVPS13A. The gene model was annotated using a hmmsearch of Pfam domains and shows conservation of VPS13 structural domains at the N-terminus and C-terminus (yellow). AtVPS13a carries a unique C2 domain (cyan) not found in other homologs.

### *AtVPS13a* is required for efficient pollen germination

To further specify the role of AtVPS13a in early pollination, we next performed reciprocal crosses between heterozygous *AtVPS13a*+/- and WT plants. We found that *AtVPS13a* is required for male gametophytic function but is dispensable for female function (Supplemental Figure S4). We confirmed that *atvps13a* pollen properly developed to mature pollen grain as two sperm nuclei and a vegetative nucleus could be observed by DAPI staining (Supplemental Figure S5).

To study the defect of *atvps13a* pollen, we used a micromanipulator to transfer pollen grains onto stigmatic papilla cells and followed their hydration and germination processes (Figure 3A). The time needed for complete hydration of the *atvps13a* pollen (mean ± SE, 8.8 ± 0.29 min) was the same with WT pollen (8.7 ± 0.38 min) (Figure 3B). However, none of the *atvps13a* pollen grains germinated within the time observed (45 min), while 94.1% of the WT pollen grains germinated at an average time of 21.4 ± 0.52 min (Figure 3C). In addition, the *atvps13a* pollen grains germinated less efficiently under *in vitro* conditions (Figure 3D). We found no pollen germination defects in a CRISPR/Cas9 (Wang et al., 2015) genome edited *atvps13b* null mutant (Figure 3C, D; Supplemental Figure S6), suggesting that this closest paralog of AtVPS13a is dispensable for its function in pollen germination. This is supported by the fact that *AtVPS13a* is expressed at much higher levels in mature anthers with pollen, than *AtVPS13b* or *SHRUBBY* (Supplemental Figure S7).

**Figure 3.**
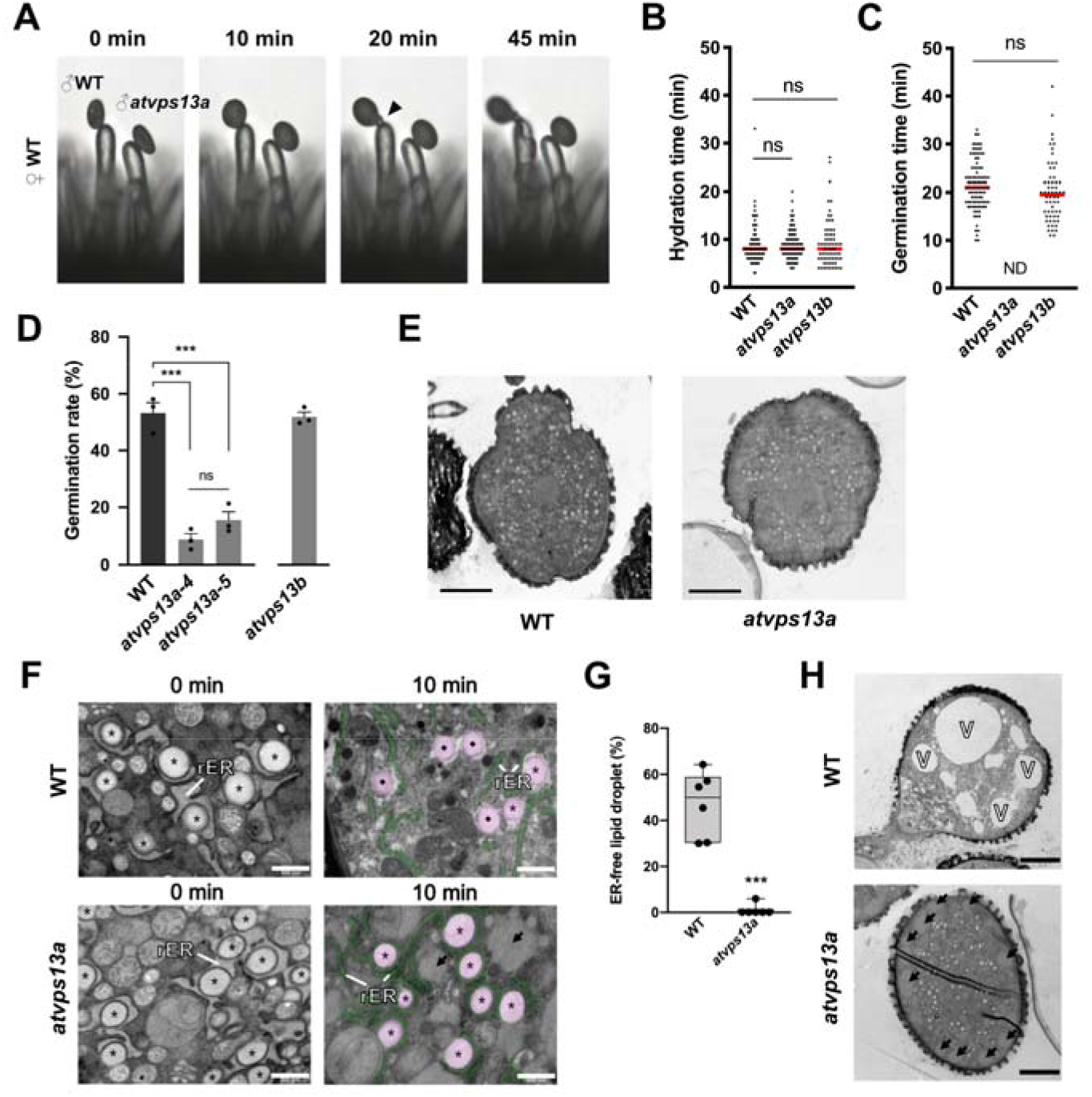
AtVPS13a is the main *Arabidopsis* VPS13 isoform required for lipid droplet-rough endoplasmic reticulum dynamic during pollen hydration. (**A**), Representative images from *in vivo* pollination assays, with the arrowhead indicating a germinated pollen tube at 20 min after pollination. (**B**), Scatter plots of pollen hydration completion times for WT, *atvps13a*, and *atvps13b* pollen on WT stigmas. Red lines, median; ns, not significant (two-tailed Student’s t-test; *p*>0.05). (**C**), Scatter plots of pollen germination times for WT, *atvps13a*, and *atvps13b* pollen on WT stigmas. Red lines, median; ND, not detected; ns, not significant (two-tailed Student’s t-test; *p* = 0.136) (B, C) WT, *n* = 102 pollen; *atvps13a, n* = 107 pollen; *atvps13b, n* = 68 pollen. (**D**), *In vitro* pollen germination rates of WT, *atvps13a-4, atvps13a-5*, and *atvps13b* pollen. Bars indicate means of replicates and whiskers indicate standard errors (*n* = 3,663, 3,135, 4930 and 3,464, respectively, where *n* is the total number of pollen grains from three replicates). Asterisks indicate significant differences by Tukey’s multiple comparisons test: \**P*<0.05, \*\**P*<0.01, \*\*\**P*<0.001; ns, not significant. (**E**), Overview of unhydrated pollen at 0 min after pollination by transmission electron microscopy (**F**), Transmission electron micrographs of unhydrated pollen at 0 min and hydrated pollen at 10 min after pollination. Pseudo color rER (green) and LD (magenta) were labeled in 10 min images to visualize LD release. (**G**), Box plots of the percentages of putative rER-free lipid droplets in hydrated pollen grains at 10 min after pollination. Asterisks indicate significant differences by two-tailed Student’s t-test: \**p*<0.05, \*\**p*<0.01, \*\*\**p*<0.001; *n* = 6 pollen grains. (**H**), Germinated WT pollen and arrested *atvsp13a* pollen at 20 min after pollination (F, H) rER, rough endoplasmic reticulum; asterisk, rER-bound lipid droplet; diamond, putative rER-free lipid droplet; arrow, putative vesicle fusion structure; V, vacuole. Scale bars: (E, H) = 5 μm, (F) = 500 nm.

### The *atvps13a* mutant failed to dissociate rER-LD and formed ectopic vesicle fusions

We next used the transmission electron microscopy to observe subcellular structures of *atvps13a* pollen after pollination. In the unhydrated phase immediately after pollination, no obvious structural defects were found in the mutant pollen (Figure 3E). rER-bound lipid droplets (LDs), also observed in a previous study (Yamamoto et al., 2003), were abundant in both WT and *atvps13a* pollen grains (Figure 3F). At 10 minutes after pollination, 47.0 ± 5.9% (mean ± SE) of the LDs had dissociated from the rER in WT pollen (Figure 3F, G), whereas the dissociation was rarely observed in the *atvps13a* mutant (1.0 ± 1.0%). At 20 minutes after pollination, the WT pollen grains formed enlarged vacuoles on the side opposite from the germination site, while such structure was not found in any of the mutant pollen (Figure 3H). Pollen grains are enriched in small golgi-derived vesicles (Yamamoto et al., 2003), and we found putative vesicle-fused structures populated near plasma membrane in the *atvps13a* mutant (‘arrow’ in Figure 3F, H). They were being surrounded by golgi-derived vesicles with similar electron-dense property, suggesting that they were formed through aberrant fusions of these vesicles (Supplemental Figure S8). These observations revealed that AtVPS13a may be required to regulate organelle reorganization during pollen germination.

### AtVPS13a polarizes to future germination site and pollen tube tip region in an actin-dependent manner

In order to observe AtVPS13a during pollen germination, we used the CRISPR/Cas9-mediated gene targeting (GT) (Miki et al., 2018) to tag fluorescent protein Venus before the stop codon of genomic *AtVPS13a* (Figure 4A). The homozygous GT-line (GT-AtVPS13a:Venus) showed unaffected pollen germination on stigmas suggesting that the fusion protein retained its function in pollen grains (Figure 4B). AtVPS13a:Venus fluorescence was evenly distributed in a mature pollen grain before pollination (Figure 4C). Live-imaging of GT-AtVPS13a:Venus pollen germination *in vitro* showed that AtVPS13a:Venus was intensified at the future germination site and later focused in elongating pollen tube tip region (Figure 4D-F). The result showed that AtVPS13a participates in cellular polarization during pollen germination.

**Figure 4.**
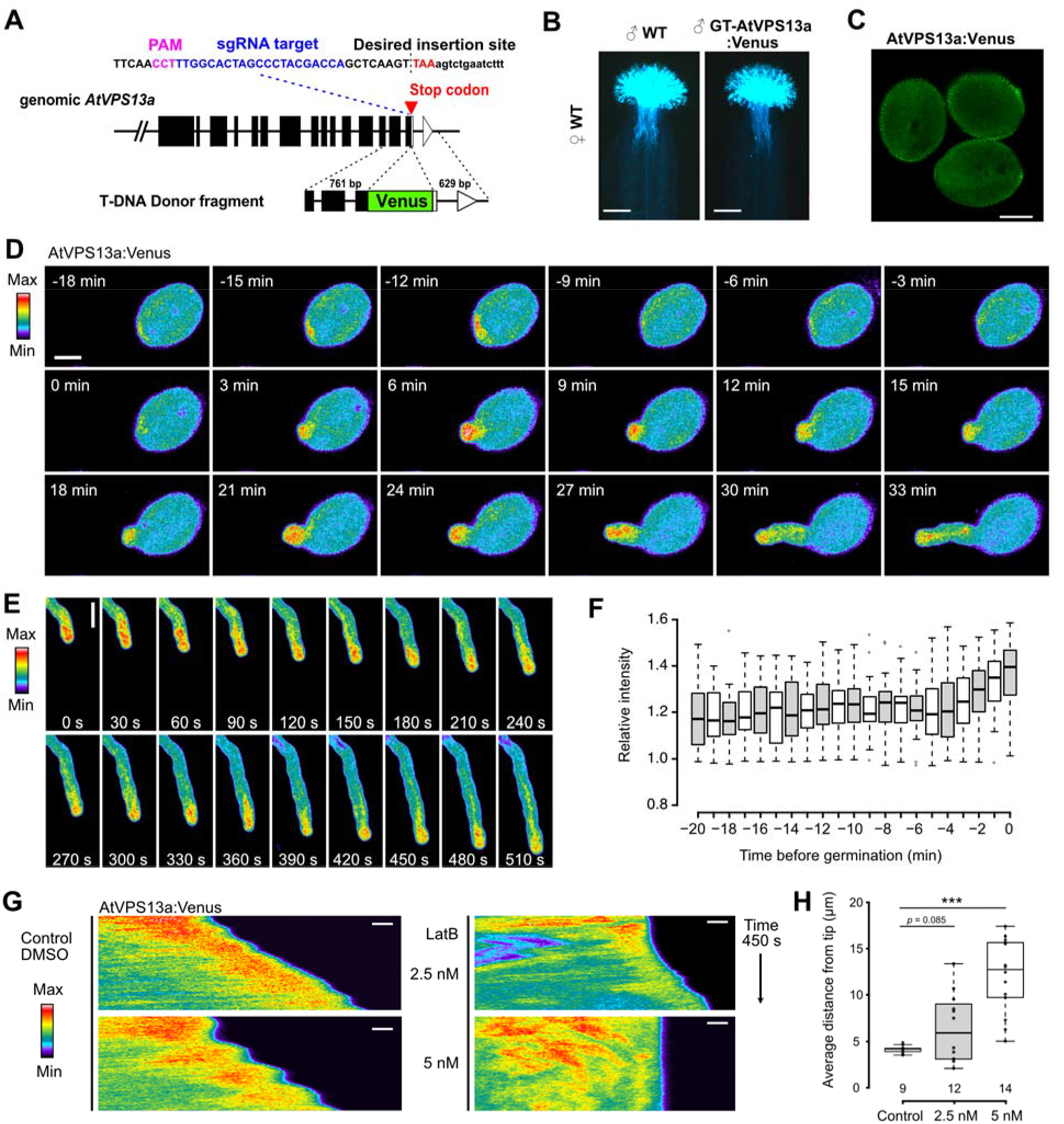
AtVPS13a:Venus polarized to future germination site and maintained at pollen tube tip by intact actin filament network. (**A**), CRISPR/Cas9 mediated gene targeting of *AtVPS13a*. The sgRNA target and PAM sequence near the stop codon are shown. A vector containing donor fragment having homology regions with desired insertion site and sgRNA expressing cassette was transferred into Cas9-expressing *Arabidopsis*. (**B**), Aniline blue staining of pistils after pollination with WT or homozygous T_3_ GT-AtVPS13a:Venus pollen for 3 h, scale bars = 200 μm. (**C**), Pollen grains from homozygous T_3_ GT-AtVPS13a:Venus plant excited by 514 nm laser. Signal from pollen exine resulted from autofluorescence of the pollen wall, scale bar = 10 μm. (**D**) and (**E**), Representative time-lapse images of *in vitro* pollen germination of GT-AtVPS13a:Venus pollen, AtVPS13a:Venus signal increased at future germination site of germinating pollen (D) and pollen tube tip of germinated pollen (D, E), scale bars = 10 μm. (**F**), Box plot of relative Venus fluorescence intensity of average signal in a circle below future germination (5μm diameter) against average signal of the whole pollen grain, *n* =19 pollen grains. (**G**), Representative kymographs of pollen tubes generated from time-lapse imaging at 3 sec intervals for 450 sec. Time-lapse was captured from 10-30 min after DMSO control or LatB treatment, scale bar = 2 μm. (**H**), Box plot of average distance of peak Venus signal from pollen tube tip, see material and method for detail, the number below each box plot represent the number of pollen tube (*n*) analyzed; *p* value calculated from two-tailed Student’s t-test; ***, *p*<0.001.

Establishment of cell polarity for pollen germination and tube growth relies on regulated arrangement of F-actin (Wu et al., 2010; Zhu et al., 2017; Liu et al., 2018). Under control condition, AtVPS13a:Venus localization focused at pollen tube tip region during both oscillatory and non-oscillatory growth with the peak of averaged fluorescence positioned at 4.17 ± 0.44 μm (mean ± SD) from pollen tube tip (Figure 4G, H). During the oscillatory growth, AtVPS13a:Venus signal appeared closer to the tip when pollen tube actively elongated (Figure 4G). The tip localization of AtVPS13a:Venus was visibly affected when 2.5 nM latrunculin B (LatB), an actin polymerization inhibitor, was treated to growing pollen tube (6.43 ± 3.68 μm); while 5 nM LatB disrupted AtVPS13a:Venus tip localization completely (12.10 ± 3.96 μm, Figure 4G, H). Effect of LatB showed that AtVPS13a:Venus localization during pollen tube tip elongation relies on correct organization of F-actin.

### AtVPS13a co-localizes with the vesicle secretion system

To gain insight on the molecular localization of AtVPS13a, we used a combination of biochemical analyses. Crude flower extract of the GT-VPS13a:Venus line was fractionated by differential centrifugation. The AtVPS13a:Venus fusion protein was most enriched in the final 100,000xg pellet which presumably contained microsomes (Figure 5A). Thus, we further fractionated the microsome fraction isolated from mature pollen of the GT-AtVPS13a:Venus line by sucrose step gradient centrifugation (Figure 5B). AtVPS13a:Venus was most enriched in the 12% sucrose fraction, but exhibited different distribution pattern from all other organelles initially investigated (Figure 5C).

**Figure 5.**
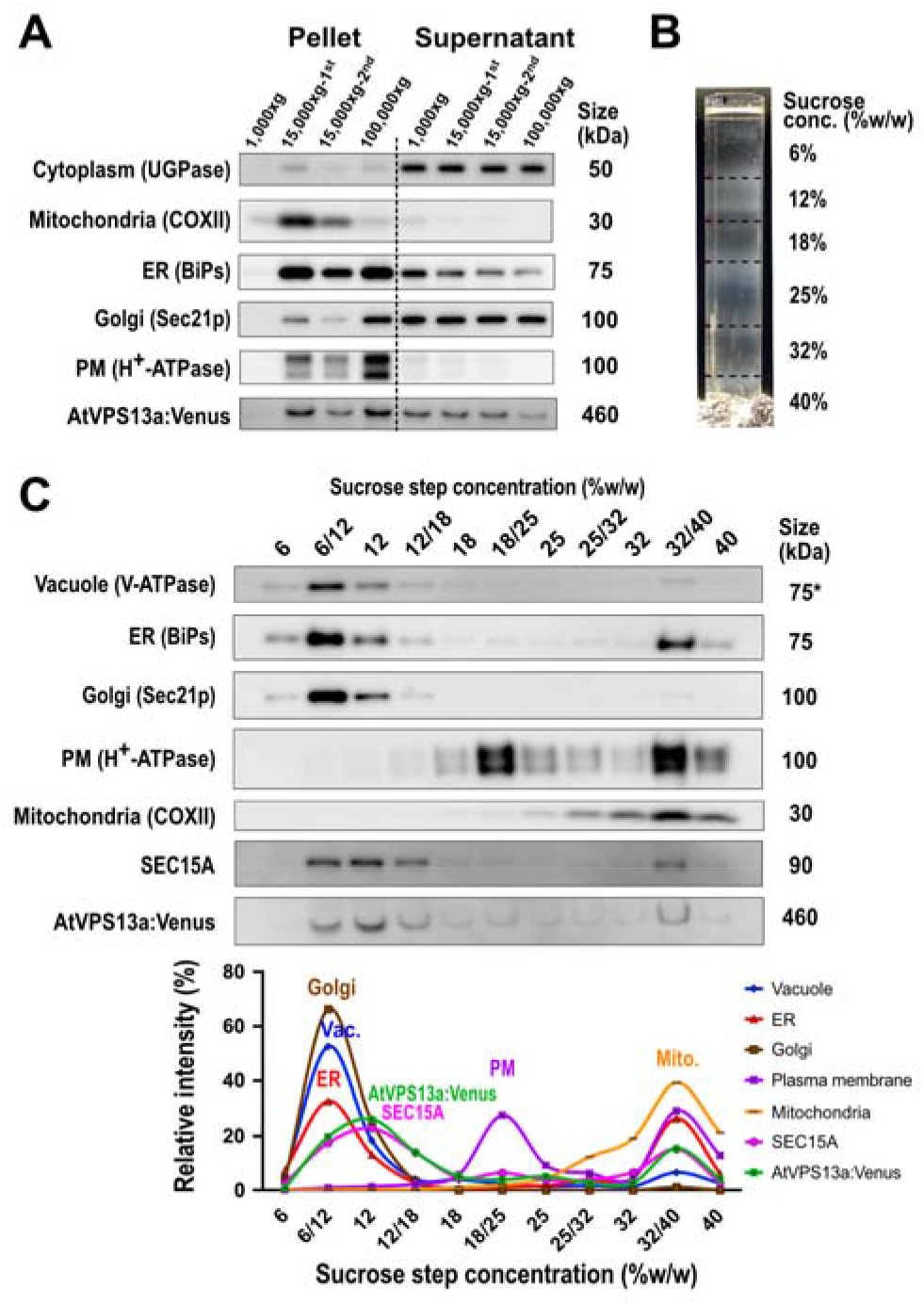
Identification of AtVPS13a:Venus coenriched proteins by proteomics. (**A**), Separation of organelles by differential centrifugation, antibodies against various organelle markers are used to visualize organelle distribution in each fraction. An antibody against GFP is used to detect AtVPS13a:Venus. The fractionation was performed independently three times using open flower samples collected on different days. The three replicates yielded similar results. (**B**), Separation of 100,000xg pellet of GT-VPS13a:Venus pollen protein by sucrose step gradient centrifugation. (**C**), Western blot analysis of each fraction from sucrose step gradient centrifugation to analyze the distribution patterns of organelle markers, AtVPS13a:Venus, and exocyst subunit SEC15A. Line graphs represent quantitative distribution of each component calculated by band intensity.

To find out cell components that co-migrated with AtVPS13a:Venus, four sucrose fractions 6/12%, 12%, 12/18%, and 18/25% were subjected to proteomic analysis by LC-MS/MS. Out of 3,046 detected proteins, distribution pattern of 340 proteins showed correlation with AtVPS13a-Venus (Pearson’s correlation coefficient >0.8, Supplemental table S1). Gene ontology (GO) enrichment analysis of these 340 proteins using PANTHER (Mi et al., 2019) identified proteins that belong to “Vesicle transport along actin filament”, “Golgi to plasma membrane transport” or “exocytosis” co-migrated with AtVPS13a:Venus (Supplemental Table S2). In agreement with the proteomic analysis, distribution pattern of an exocyst subunit SEC15A investigated by its specific antibody strongly resembled that of AtVPS13a:Venus (Figure 5C). We also performed co-immunopurification (co-IP) of crude pollen proteins from the GT-AtVPS13a:Venus line. In total, 82 proteins were uniquely detected in the AtVPS13a:Venus co-IP fraction but were not detected in that of WT (Supplemental Table S3). GO analysis showed that “exocytosis” was the most significantly enriched term (*P* = 3.32e-03) in the 82 proteins. Notably, exocyst complex subunits SEC8 and SEC5B that are required for exocytosis during pollen germination (Hála et al., 2008), and Armadillo Repeat Only1 (ARO1) involved in polar cell growth (Gebert et al., 2008) co-localized with AtVPS13a in both sucrose gradient purification and co-IP (Supplemental Table S1, S3).

### AtVPS13a synchronizes with vesicle polarization following local Ca^2+^ signal

During progression towards pollen germination, polarized vesicle secretion and Ca^2+^ gradient formation at the future germination site occur (Iwano et al., 2004; Hoedemaekers et al., 2015), and from the above section we found that AtVPS13a co-localize with the vesicle-related systems. Thus, we investigated if AtVPS13a distribution dynamics correlates with secretory vesicle movement or Ca^2+^ gradient during pollen germination. Secretory vesicle marker mCherry:RabA4B or Ca^2+^ reporter R-GECO1 were introduced into the GT-AtVPS13a:Venus plant background to generate dual reporter lines. Interestingly, simultaneous live imaging of secretory vesicles and AtVPS13a:Venus during *in vitro* germination showed that both signals accumulated at the polarized site with similar timing (Figure 6A-C; Supplemental Figure S9; Supplemental Movie S1). In contrast, each local peak of AtVPS13a:Venus lagged 3.03 ± 0.529 min behind a Ca^2+^ peak at a polarized site (Figure 6C-E, Supplemental Figure S9). These results showed that AtVPS13a and RabA4B vesicle marker had synchronized local accumulation after a Ca^2+^ elevation at future pollen germination site.

**Figure 6.**
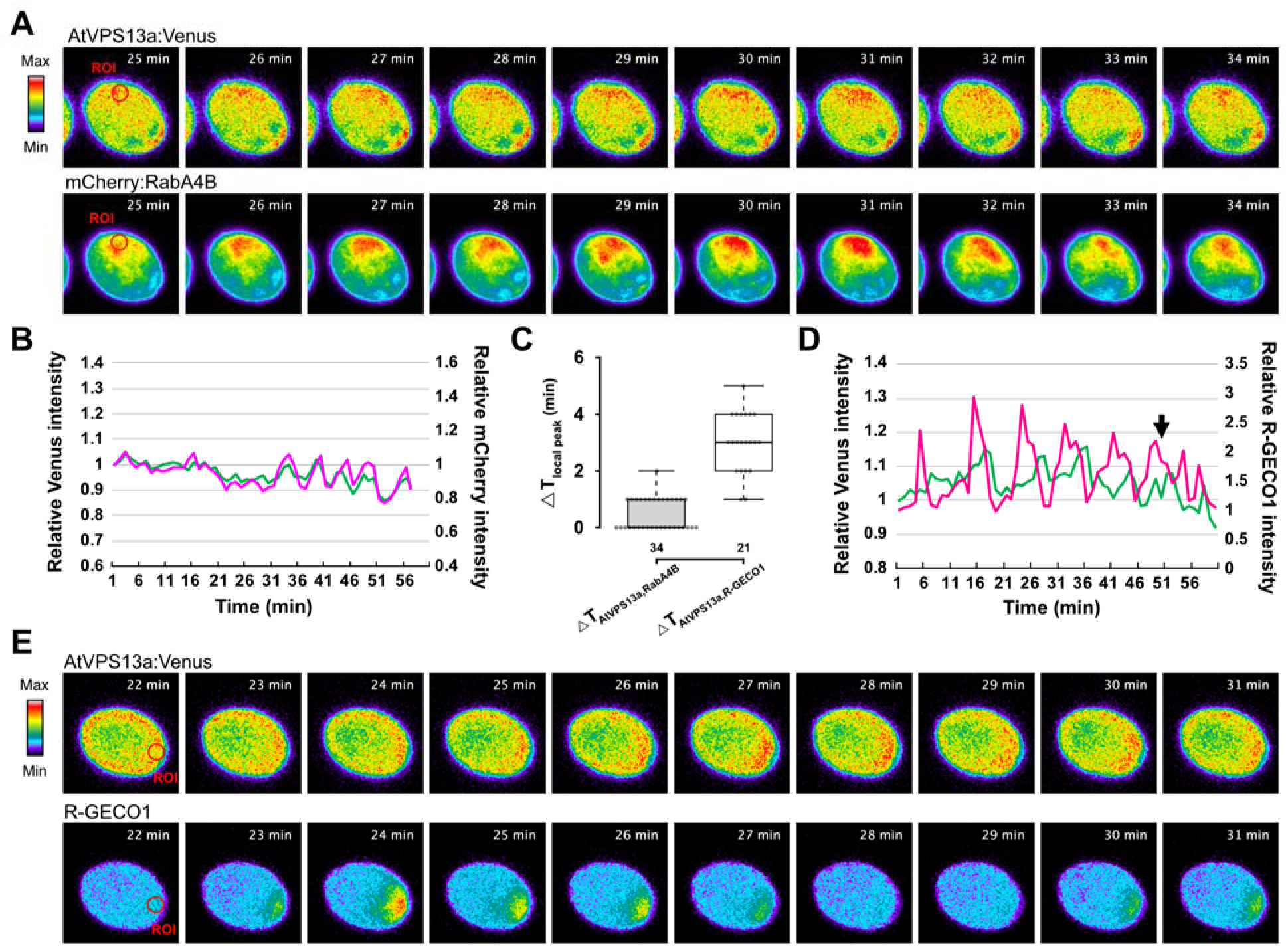
Ca^2+^ signaling preceded AtVPS13a:Venus and mCherry:RabA4B vesicle polarization in activated pollen grains. (**A**), Representative time-lapse images of *in vitro* pollen germination of the dual reporter, AtVPS13a:Venus and mCherry:RabA4B pollen; ROIs indicate regions of interest for relative intensity calculation in (B). (**B)**, Dynamic of relative intensity of AtVPS13a:Venus (green) and mCherry:RabA4B (magenta) at the polarized site (ROI). (**C**), Time difference between each oscillating fluorescence peak of AtVPS13a and RabA4B or R-GECO1 during *in vitro* polarization. (**D**), Dynamic of relative intensity of AtVPS13a:Venus (green) and R-GECO1 (magenta-red) at the polarized site (ROI), arrow indicates the germination of the pollen grain. (**E**), Representative time-lapse images of *in vitro* pollen germination of the dual reporter, AtVPS13a:Venus and R-GECO1 pollen grain.

### AtVPS13a mediates vesicle polarization downstream of Ca^2+^ signaling

It has been known that Ca^2+^ gradient forms within pollen grain, most concentrated at the papilla cell contact site after pollen hydration (Iwano et al., 2004). We investigated the local Ca^2+^ increase inside the pollen grain by the Ca^2+^ sensor, Yellow Cameleon 3.6 (YC3.6), after pollination. In agreement with the previous report, cytoplasmic Ca^2+^ level rapidly decreased during pollen hydration (Iwano et al., 2004) (Figure 7A). We further found that the first Ca^2+^ spike appeared only briefly (shorter than 30 sec, Supplemental Movie S2) at 4.87 ± 1.70 min (mean ± SD) after hydration initiation in the WT (Figure 7B). The *atvps13a*/YC3.6 pollen also showed the Ca^2+^ spike at the same timing (5.10 ± 1.69 min; two-tailed Student’s t-test, *p* = 0.559). After the dissipation of the first Ca^2+^ spike, gradual cytoplasmic Ca^2+^ increase until germination was observed (Figure 7C, Supplemental Movie S2). This Ca^2+^ dynamic was similarly observed in both WT and *atvps13a* pollen grains (Figure 7A-C), suggesting that AtVPS13a is not required to trigger this signal, but functions to establish pollen polarity downstream of the Ca^2+^ gradient.

**Figure 7.**
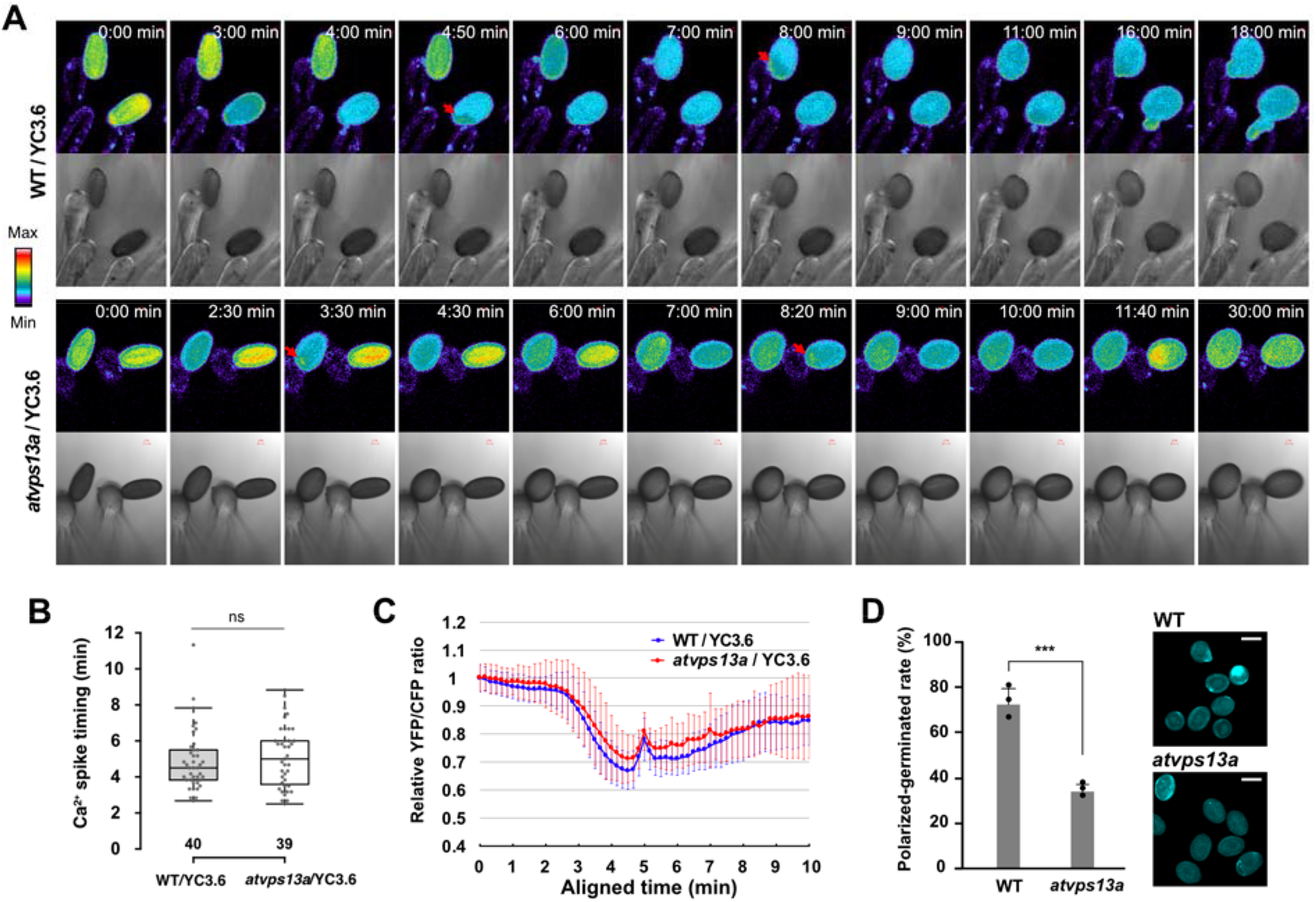
AtVPS13a mediates pollen polarization downstream of Ca^2+^ signaling. (**A**), Ratiometric images of adjusted YFP/CFP ratio from YC3.6 representing Ca^2+^ level in WT and *atvps13a* plant background. Red arrows indicate the first Ca^2+^ spike during pollen hydration of both WT and mutant pollen grains. (**B**), Box plot of the first Ca^2+^ spike timing after pollen hydration initiation. The number below each box plot represent the number of pollen grain observed (*n*); ns indicated non-significant difference between means of the two groups by two-tailed Student’s t-test (*p* = 0.559). (**C**), Pollen Ca^2+^ dynamic at the papilla cell contact represented by ratiometric images of YFP/CFP of YC3.6. Time after pollination was aligned in the manner that makes the first Ca^2+^ spike appear at 5 min. (**D**), Sum of polarized pollen (marked by callose deposition at pollen wall) and germinated pollen rate after incubation in liquid pollen germination medium and stained with aniline blue. Asterisks indicate significant difference by two-tailed Student’s t-test: *** *p*<0.001; scale bars = 20 μm.

AtVPS13a has a predicted Ca^2+^-dependent lipid-binding C2 domain (Fig 2B) which may mediate Ca^2+^ signaling response during pollen germination. We used CRISPR/Cas9 to generate the *AtVPS13a* mutant lacking complete C2 domain and found that pollen tube tip focusing of AtVPS13aΔC2:Venus was partially lost (Supplemental Figure S10A-C). The result indicates that C2 domain region was required to regulate the cellular distribution of AtVPS13a. Pollen germination rate and tube growth were also affected in the C2 deletion lines (Supplemental Figure S10B, D). This may have resulted from the combined effect of the reduced polarity and protein stability, since *AtVPS13aΔC2:Venus* pollen accumulated reduced amount of the fusion protein (Supplemental Figure S10E).

Considering that AtVPS13a co-localized with secretory vesicles in the polarizing pollen (Figure 6A, B), we speculated that AtVPS13a supports cell wall secretion for successful germination. We incubated pollen grains in liquid pollen germination medium and used aniline blue staining to observe callose deposition onto pollen cell wall. We observed at least one callose spot in 72.6 ± 6.90 % (mean ± SD) of the WT pollen but only 33.8 ± 2.90 % in *atvps13a* pollen (Figure 7D). Despite sensing the same Ca^2+^ cue like WT *in vivo* (Figure 7A-C), the result strongly suggests that *atvps13a* pollen has limited vesicle secretion at the potential germination site, thus unable to deliver enough cell wall carbohydrate for rapid germination.

## DISCUSSION

In this study, we rediscovered an overlooked phenomenon as an essential step toward successful pollen germination. In *Brassica oleracea*, LDs were found unwinded from ER in hydrating pollen grains (Elleman and Dickinson, 1986). We found that this process was mediated by AtVPS13a in *Arabidopsis* pollen because LDs in hydrated *atvps13a* pollen remained tightly bound to rER (Figure 3F, G). We speculate that AtVPS13a-mediated LD release is required to expose its surface for further catabolism. There are some reports on active consumption (Zienkiewicz et al., 2010, 2012) and structural protein degradation (Kretzschmar et al., 2018) of LDs in pollen. It is possible that AtVPS13a-mediated LD release is a prerequisite which primes pollen grain for pollen polar growth.

Since we found that AtVPS13a biochemically co-localized with the exocyst subunits (SEC5B and SEC8), our study showed that AtVPS13a may be functionally related to exocytosis. It is known that the exocyst complex comprises subcomplexes which dynamically bind cytoplasmic secretory vesicle (Ahmed et al., 2018). The secretory vesicle is then tethered to the destined membrane by SEC3 and EXO70 subunits before subsequent membrane fusion (Ahmed et al., 2018; Mei and Guo, 2018). We found that AtVPS13a and RabA4B secretory vesicle marker synchronized their accumulation at the germination site (Figure 6A-C), where exocytosis takes place to mediate cell wall deposition (Hála et al., 2008; Bloch et al., 2016; Li et al., 2017). Furthermore, *atvps13a* pollen formed exocytotic vesicle fusion structures in hydrated pollen grains (Supplemental Figure S8), indicating that AtVPS13a is required to prevent ectopic fusion of these vesicles. We also found that AtVPS13a co-localize with a member of the ARO subfamily proteins, recently reported to have a scaffolding function to limit ROP signaling to the polar growth sites (Kulich et al., 2020). Pollen ARO1, biochemically co-localized with AtVPS13a (Supplemental Table S1, S3), localizes to future germination site (Vogler et al., 2015), and is required for tip growth maintenance by regulating F-actin network (Gebert et al., 2008). It is possible that AtVPS13a is needed to establish polarized exocytosis by co-operating with the exocyst complex and/or ARO1.

After pollination, pollen uptake water and extracellular Ca^2+^ through the contact with the papilla cell, which may provide directional cues for the pollen tube emergence site (Lush et al., 1998; Iwano et al., 2004; Swanson et al., 2004). We discovered that pollen briefly uptake extracellular Ca^2+^ from the papilla contact site before hydration completion (Figure 7A, B). To our knowledge, this study is the first to visualize this brief Ca^2+^ uptake as early as 5 min after hydration initiation, and AtVPS13a and RabA4B may be recruited to this Ca^2+^ elevated site (Figure 6A-E). Intriguingly, C2 domain was required for efficient polarized localization of AtVPS13a (Supplemental Figure S10B, C). Considering the Ca^2+^-dependent membrane localization of a C2 domain (Corbalan-Garcia and Gómez-Fernández, 2014), AtVPS13a may mediate pollen polarization downstream of local Ca^2+^ signal to ensure proper vesicle trafficking to the germination site.

Recent studies found that VPS13 proteins tether or transport lipid between organelles (Li et al., 2020; Cai et al., 2022). They localize to various organelle contact sites depending on cell type and cellular context (Park and Neiman, 2012; Kumar et al., 2018; Kolakowski et al., 2021; Wang et al., 2021). AtVPS13a contained hydrophobic cavity at its N-terminus that may transfer lipids (Li et al., 2020; Cai et al., 2022), and also the amphipathic helix at its C-terminus that may be anchored to a biological membrane (Kumar et al., 2018) (Supplemental Figure S3A, B) like other VPS13s. Although it remains speculative, it is conceivable that AtVPS13a may mobilize lipid from a source such as free LD to other organelles either directly or through intermediate membrane transport system such as vesicles. Potential sink organelles are extending membranes such as vacuole or rapidly expanding plasma membrane.

## METHODS

### Plant growth conditions

Surface-sterilized seeds were sown on half-strength Murashige and Skoog medium containing 0.8% agar, stratified at 4°C in the dark for 3 days, then transferred to a growth chamber set at 22°C with a 14-h-light/10-h-dark cycle. About one-week-old seedlings were transferred and grown on soil in a growth room at 22°C with the 14-h-light/10-h-dark cycle.

### Reporter plants

All oligonucleotides used in this study are listed in Supplemental Table S4. The promoter region of At4g24570 (approximately 1 kb) was amplified from genomic *A. thaliana* Col-0 DNA. The purified fragment was cloned into pCambia1300, described in a previous study (Fujii et al., 2019), using the In-Fusion HD Cloning Kit (Takara Bio) after *Hin*dIII/*Bam*HI double digestion of the vector. The *Photinus pyralis* luciferase gene fragment was then cloned into the *Bam*HI/*Sac*I site of the vector to achieve the fusion of the At4g24570 promoter to the luciferase reporter. The construct was introduced into the Col-0 strain using the *Agrobacterium*-mediated transformation (Clough and Bent, 1998). Creation of the pollen calcium reporter line ACT1_pro_:*YC3.6* was reported previously (Iwano et al., 2009). The reporter line was crossed with *atvps13a* and the progenies were screened by PCR and fluorescence microscopy to obtain homozygous *atvps13a* individuals expressing YC3.6 (*atvps13a*/ACT1_pro_:*YC3.6*). To generate dual reporter lines; UBQ10_pro_, mCherry::HSP_ter_, and RabA4B were sequentially cloned into pCAMBIA1300 vector using In-Fusion cloning technology (Takara Bio) to construct UBQ10_pro_:mCherry:RabA4B:HSP_ter_ cassette. For the R-GECO1 construct, R-GECO1:HSP_ter_ was cloned after UBQ10_pro_ in pCAMBIA1300 backbone. *A. tumefaciens* containing each plasmid was then used to transform GT-VPS13a:Venus plant.

### T-DNA lines and mutant plants

T-DNA lines were obtained from the Arabidopsis Biological Resource Center (ABRC). Their stock codes for the lines used in this study are as follows: *cer1*, SALK_008544C; *cer6-2*, CS6252; *fkp1*, SAIL_675_A04; *atvps13a-5*, SALK_058035. For the genetic screen, plants grown from ethyl methanesulfonate (EMS)-treated seeds were referred to as the M_1_ generation. Selfed-seed from each M_1_ individual was collected separately, and about five M_2_ plants from each M_1_ individual were sown on soil. Pollen from approximately 2,400 M_2_ individuals was screened for pollen activity using the luciferase-based pollination assay described below. The *atvps13a-1–4* mutant alleles were generated in this EMS mutagenesis procedure. The *atvps13a-4* plants used for further luciferase-based pollination assays and *in vitro* germination assays were BC_1_F_2_ and BC_1_F_3_ progeny derived from a cross between an EMS M_2_ mutant and Col-0.

### Luciferase-based pollination assay

Flower buds of reporter plants were emasculated one day before the assay. On the day of the assay, a LUMITRAC™200 microplate (Greiner Bio-One) was prepared by filling each well with 280 μl 1% agar, then a pollen-free reporter pistil was stood in each well. A small drop of 1 mM D-luciferin + 0.025% Tween 20 was pipetted onto the reporter stigma and allowed to air dry at room temperature for 1 h. The pistil was then pollinated with the pollen of interest. Bioluminescence was counted for 3 sec/well at 3 hours after pollination for end-point measurements, or every 3 minutes for time course measurements, using a TriStar^2^ LB942 Modular Multimode Microplate Reader (Berthold). The relative increase in bioluminescence was calculated in relation to the bioluminescence of unpollinated reporter pistils.

### Genome deep sequencing for causal mutation detection

F_1_ plants were obtained from the crosses between the EMS M_2_ mutants and Col-0. F_2_ plants were screened for their lack of ability to induce the pistil bioluminescence of the reporter plants. Pooled genomic DNA samples from about 20 individual homozygous recessive mutants were extracted using the DNeasy Plant Mini Kit (Qiagen) and concentrated by ethanol precipitation. The sequencing analysis and identification of mutations was performed as described in ref. (Suzuki et al., 2018).

### Aniline blue staining

Pollinated pistils were fixed in ethanol:acetic acid (3:1) overnight. Fixed tissues were incubated in 1 N NaOH at 60°C for 30 minutes before staining in 2% K_3_PO_4_ + 0.01 % aniline blue solution in the dark for at least 3 hours at room temperature, or for extended periods at 4 °C. The specimens were mounted in 50% glycerol and observed using an epifluorescence microscope with the Zeiss filter set 01 (excitation BP 365/12, Emission LP 397). To stain pollen sample after *in vitro* germination, pollen was pelleted after incubation in liquid pollen germination medium and resuspended in 2% K_3_PO_4_ + 0.01 % aniline blue solution to stain callose before observation by confocal microscope, LSM880 (Zeiss).

### Phylogenetic analysis

Animal, plant, and yeast proteomes were obtained from Ensembl (ftp://ftp.hgc.jp/pub/mirror/ensembl/current_fasta/), the DOE Joint Genome Institute (https://genome.jgi.doe.gov/portal/pages/dynamicOrganismDownload.jsf?organism=Phytozome), and the Saccaromyces Genome Database (https://www.yeastgenome.org/), respectively (Supplemental table S5). HMM profiles of Chorein_N (PF12624), VPS13 (PF16908), SHR-BD (PF06650), and VPS13_C (PF16909) were obtained from Pfam (https://pfam.xfam.org/). We searched each proteome for each of these domains using hmmscan, which is included in the HMMER package (version 3.2.1; http://hmmer.org/). The threshold for E-value and domE-value were 1E-05. In this study, we defined “VPS13” as the proteins possessing all four domains mentioned above. For construction of the VPS13 phylogenetic tree, sequences of the four domains were extracted from the VPS13 proteins. For each protein, only the domain with the lowest domE-value from the hmmscan was used when multiple sites were found. Multiple alignments of each domain were done using the Clustal Omega program version 1.2.4 (Sievers et al., 2011). All alignments were combined, and gap regions were removed manually. A Bayesian inference was performed using MrBayes version 3.2.6 (Ronquist et al., 2012). The Metropolis coupled Markov Chain Monte Carlo processes were run for 500,000 generations, and trees were collected every 100 generations. After discarding trees corresponding to the first 25% (burn-in), the remaining trees were used to generate the consensus phylogenetic tree. Bayesian posterior probabilities were estimated as the proportion of trees sampled after burn-in.

### Reciprocal cross experiment

Reciprocal crosses were performed between a WT Col-0 plant and a heterozygous SALK_058035 *AtVPS13a*+/-mutant plant carrying the Kanamycin resistance gene in the middle of the *AtVPS13a* gene (the *atvps13a-5* allele). Immature flowers of each plant were emasculated before anthesis and pollinated with pollen from the other genotype. The experiment was repeated each day on the same plant pair for about one week. Collected seeds were screened on half-strength Murashige and Skoog medium supplemented with 50 μg/ml kanamycin, and the seedlings were phenotyped. A Chi-square test was used to compare segregation of the *AtVPS13a* alleles with the expected distribution by Mendelian’s law (1:1 ratio of Km_r_:Km_s_).

### *In vivo* pollination assay

To determine the time pollen grains needed to fully hydrate and germinate on stigmas, pollination assays were performed as previously described (Iwano et al., 2014), with a slight modification The un-pollinated WT pistils were fixed with double-sided tape to glass slides and kept hydrated in 1% agar. The slides were set on an Axiovert 135 (Zeiss) inverted microscope equipped with a micromanipulator, which was used to place individual pollen grains on different papilla cells (one grain per cell). Photos were taken every minute for 45 min for the following numbers of individual pollen grains: *n*; WT = 102; *atvps13a* = 107; *atvps13b* = 68 pollination events.

### *In vitro* pollen germination

*In vitro* pollen germination was performed as described previously (Li et al., 1999) with slight modifications. The pollen germination medium (PGM) contained 18% sucrose, 1 mM CaCl_2_, 1 mM Ca(NO_3_)_2_, 1 mM MgSO_4_, 0.01% Boric acid, 1 mM PIPES pH 7.0, and 1% Nusieve™ GTG™ Agarose (Lonza) in ultrapure water. Pollen was carefully transferred from 3–5 open flowers onto a 1 mm-thick pollen germination medium pad. Pollen of different genotypes were germinated on the same germination pad in each replicate. The germination pad was incubated in a humid chamber at 22°C for one day before observation. Replicates with Col-0, *atvps13a-4*, and *atvps13a-5* pollen were performed on three different days. The results for *atvps13b* pollen were from germination on three germination pads in one day. ANOVA was used to compare the mean germination rates. For live-imaging and callose deposition experiments, liquid PGM (agarose omitted) supplemented with 10 μM Epibrassinolide (epiBL; E1641, Sigma) was used to increase pollen germination efficiency (Vogler et al., 2014). About 15 flowers were collected into 1.5 ml tube and vigorously vortexed in 500 μl PGM+epiBL for 1 min to separated pollen grain from dehisced anther. The tube was incubated at room temperature for 5 min and centrifuged at 500xg 2 min to pellet pollen grain. Separated pollen pellet was resuspended in a fresh 100-150 μl PGM+epiBL and transfer onto glass bottom dish for observation under confocal microscope.

### Confocal imaging and Latrunculin B treatment

All imaging was performed using Zeiss LSM 880 confocal laser scanning microscope (Zeiss) controlled by ZEN 2.3 SP1 FP1 software (v. 14.08.201, Zeiss).

Live-imaging during *in vitro* pollen germination was acquired from pollen/pollen tube in liquid PGM+epiBL. Wide-field imaging to capture pollen grains polarization and initial germination was imaged by C-Apochromat 40x/1.1 W Corr M27 water immersion lens at 1 min intervals for 60 cycles. Pollen tube was imaged by Plan-Apochromat 63x/1.4 Oil DIC M27 immersion lens at 3 sec intervals for 150 cycles. For AtVPS13a:Venus dynamic during pollen germination, Venus was excited by 514 nm Argon laser and emission (em.) wavelength was captured at 517-588 nm. In dual-labeled lines, fluorescent protein variants were excited and acquired simultaneously (switch track every line); Venus was excited by 488 nm Argon laser (em. 517-562 nm), while mCherry and R-GECO1 was excited by 561 nm DPSS laser (em. 602-660 nm).

For imaging of pollen tube after Latrunculin B (LatB; L5288, Sigma) treatment, LatB stock solution in DMSO was first diluted to 2x working concentration in liquid PGM+epiBL, PGM+2xLatB was then added to germinated pollen solution on glass bottom dish at 1:1 ratio. Pollen tube was imaged after 10-30 min incubation (2-3 pollen tubes/one experiment). DMSO treated pollen tube was used as control.

To observe pollen Ca^2+^ dynamic after *in vivo* pollination. Unpollinated pistils were prepared as described above for the *in vivo* pollination assay, however, set up on LSM 880. After transferring pollen grain by micromanipulator, YC3.6 FRET was observed using Plan-Apochromat 20x/0.8 M27 objective lens. The YC3.6 protein was excited with a 440 nm Diode laser, and fluorescent emissions from ECFP (CFP) and cpVenus (YFP, from FRET) were collected at 464–499 nm and 526–553 nm, respectively. The fluorescence was collected every 10 s for 30–50 min.

To observe callose deposition on pollen cell wall, pollen incubated in liquid PGM+epiBL was directly stained by aniline blue solution. Plan-Apochromat 20x/0.8 M27 objective lens was used to capture wide-field 3*3 tile scanning (10% overlap at tile boundary). Stained pollen was excited by 440 nm diode laser and emission wavelength was captured at 445-589 nm. Intensities from three Z-stacks with 3 μm intervals were projected onto single 2D image by ZEN 2.3 software before scoring callose spot.

### Image processing and quantification

To measure AtVPS13a:Venus dynamic at future pollen germination site, a circular region of interest (ROI) with 5 μm diameter beneath the pollen tube emergence site and ellipsoid ROI covering whole pollen grain were drawn. Average intensity value of each ROI of every time point was exported from ZEN 2.3 software. Intensity at germination site ROI was divided by whole grain ROI for normalization, and the relative intensity was plotted against time to show AtVPS13a:Venus dynamic at future germination site. Similar measurement was performed for dual-labeled pollen, AtVPS13a:Venus with mCherry:RabA4b or R-GECO1, but the signal intensity was normalized by intensity of the first time point rather than the whole grain ROI.

For measurement of average distance between AtVPS13a:Venus peak signal from pollen tube tip, kymograph of growing pollen tube was analyzed by Fiji software (Schindelin et al., 2012). Moving average of 3*3 pixels was applied to raw time-lapse images to reduce noise. Kymograph was generated by intensity profile of the line along the middle of each pollen tube (line width = 10) using ZEN 2.3 software. One horizontal pixel row in a kymograph image represents one image of pollen tube (150 time-lapse cycles generated 150 pixels kymograph). Exported grey-scale kymograph was imported into Fiji and 10 pixels-height rectangle ROI was drawn to extract intensity profile from the kymograph (kymograph binning, 1 rectangle ROI profile equals to averaged profile of 10 consecutive pollen tube images). The kymograph profile extraction was repeated as the rectangle ROI moved through the kymograph along the Y axis at 5 pixels/step creating a total of 29 overlapping bins of averaged Venus signals along the middle of the pollen tube. The data matrix was handled in Microsoft Excel to identify the distance between the point of highest averaged Venus intensity and the tip of the pollen tube. Average distance from 29 bins was used as a “Average distance from tip” of each pollen tube.

For measurement of *in vivo* Ca^2+^ dynamic in YC3.6 reporter lines, moving average of 4*4 pixels was applied to raw time-lapse images to reduce noise. Image calculator processing by Zen 2.3 software was performed with the equation (YFP/CFP)*4000 to get YFP/CFP ratiometric images. Intensity was scaled to min/max scale of the first image, while maintaining a linear relationship, for the ease of identifying Ca^2+^ spikes. A circular ROI with 5 μm diameter was drawn beneath the pollen tube emergence site, and ROI intensity was normalized by the value of the first time point. The Ca^2+^ spike timing was defined by the time in which the relative YFP/CFP ratio reached the first peak after rapid drop during early hydration stage. The Ca^2+^ spike timing was used as a reference point of time to compare the Ca^2+^ dynamic of different pollen grains.

### CRISPR/Cas9 edited plants

CRISPR/Cas9 mediated genome editing was used to generate the *atvps13b* null mutant and the *AtVPS13aΔC2* line. Single guide RNAs (sgRNAs) were designed using the CRISPR-P 2.0 web-based service (Liu et al., 2017) (http://crispr.hzau.edu.cn). To create a frameshift in *AtVPS13b*, two sgRNA fragments targeting two genomic regions, 5’-GTTTGTTGAGAGGTCGGTAC<u>AGG</u>-3’ and 5’-GTGCTTCAAAACTTTATGAT<u>GGG</u>-3’ (protospacer adjacent motifs, PAMs, are underlined) were cloned into pHEE401E to construct an egg-cell specific expression system (following the procedure in ref. (Wang et al., 2015)). To create the in-frame deletion in *AtVPS13aΔC2*, two sgRNA fragments targeting two genomic regions, 5’-GACTATCAACAACGAGGCGT<u>AGG</u>-3’ and 5’-GGGTCACTTTCGTTTCCTGT<u>TGG</u>-3’ (PAMs are underlined), were cloned into pHEE401E. Agrobacterium-mediated transformation of either *Arabidopsis* Col-0 was done using the floral dip method (Clough and Bent, 1998). Hygromycin resistant transformants were screened for gene editing by Sanger’s sequencing.

Gene targeting (GT)-AtVPS13a:Venus plant was generated by sequential transformation method (with modification from ref. (Miki et al., 2018)). We generated an in-house vector by cloning sgRNA-expressing fragment from pHEE401E into pGWB backbone containing BASTA resistance gene. The donor fragment, AtVPS13a-donorFront:Venus:donorBack (Figure 4A) generated by overlap extension PCR, and sgRNA sequence were then sequentially cloned into the backbone. Finally, PAM sequence in the donor fragment was mutated by site-directed mutagenesis, to avoid being cleaved by Cas9 (synonymous mutation). Agrobacterium was used to introduce T-DNA from the constructed plasmid into Cas9-expressing CS69955 background (obtained from the ABRC). BASTA resistant transformants were screened for GT allele by the primer pairs which anneal genomic region but not the donor fragment within the T-DNA.

### Transmission electron microscopy

Pollinated pistils left at room temperature for 0, 10, and 20 min were cut above the style and immediately frozen in a high-pressure freezer (BAL-TEC, HPM010). The samples were fixed in 2% osmium tetroxide in anhydrous acetone and kept at low temperatures in a Cryoporter (CS-80CP, Scinics). The temperature was maintained at -80°C for 76 h, then gradually increase at a rate of 2°C/h, with holds when reaching -60°C (4 h hold), -30°C (3 h hold), and -20°C (4 h hold). The temperature was then increased to 4°C at 12°C/h and held for 2 h before transfer to room temperature. The specimen was washed with acetone and embedded in Spurr resin (Spurr Low Viscosity Embedding Kit, Polysciences). Ultra-thin sections (70–80 nm) were prepared using an ultramicrotome (Leica Ultracut UCT). The sections were stained in 4% uranyl acetate and lead citrate, then observed under a transmission electron microscope at 100 kV (JEM-1010, JEOL).

### Organelle fractionation

Every sample handling step was performed on ice. At least 60 open flowers randomly picked from five T_2_ GT-VPS13a:Venus plants were pooled and homogenized by plastic pestle in the homogenization buffer (0.5 M sucrose, 5 mM EDTA, 0.1% bovine serum albumin, 5 mM 2-mercaptoethanol, Sigma protease inhibitor cocktail, 50 mM Tris-HCl pH7.5). The homogenate was filtered through 41 μm and 11 μm nylon filters, then passed through a 5 μm Omnipore membrane filter (JMWP09025, Merck millipore). The resulting filtrate was subjected to differential centrifugation with increasing speeds; 1,000xg, 10 min; 15,000xg, 30 min; 15,000xg, 30 min; and 100,000xg, 90 min. Supernatant and pellet from each centrifugation step were kept on ice until used for SDS-PAGE and western blot analysis.

For sucrose step gradient fractionation, 6.11 g of homozygous T_3_ GT-AtVPS13a:Venus open flower was aliquoted into four 50 ml tubes and vortexed in 0.5 M sucrose 25 mM Tris-HCl pH 7.5 to isolate mature pollen. The solution was centrifuged at 500xg 2 min 4°C, pollen pellet was transferred into two 1.5 ml tubes and pelleted by centrifugation again. Pollen pellet in each tube was homogenized by pestle for 1.5 ml tube in 700 μl homogenization buffer; 0.5 M Sucrose, 5 mM EDTA, 0.1% (w/v) bovine serum albumin, 10 mM DTT, proteinase inhibitor cocktail, 50 mM Tris-HCl pH 7.5. Crude pollen extract was subjected to differential centrifugation program as stated above and supernatants were pooled before ultracentrifugation step. The microsome pellet was resuspended in 535 μl resuspension buffer (6% w/v sucrose, 1 mM EDTA, 2mM DTT, and 50 mM Tris-HCl pH 7.5) for use in sucrose step gradient centrifugation. Sucrose step gradient (sucrose in resuspension buffer) were prepared in Ultra-Clear™ centrifuge tube 11×89 mm (Beckman Coulter) starting from the bottom (%w/w); 40%, 1.5 ml; 32%, 2.2 ml; 25%, 2.2 ml; 18%, 2.2 ml; 12%, 1.5 ml; 6%, 1.5 ml. The pollen microsome fraction was gently loaded onto 6% sucrose fraction and covered on top by resuspension buffer without sucrose. The gradient was centrifuged at 100,000xg in SW 41 Ti swinging-bucket rotor (Beckman Coulter) for 2 h and each sucrose fraction and interphase were separately collected for SDS-PAGE and western blot analysis.

### Western blotting

Protein samples were run on SDS-PAGE and transfer onto PVDF membrane (Immobilon-P, Millipore). The membrane was blocked by PVDF blocking reagent (Toyobo) and rinsed by TBS-T buffer before incubated with primary and secondary antibodies, respectively. The primary antibodies used were anti-UGPase (AS05 086, Agrisera), anti-COXII (AS04 053A, Agrisera), anti-BiP (AS09 481, Agrisera), anti-Sec21p (AS08 327, Agrisera), anti-H^+^-ATPase (AS07 260, Agrisera), anti-SEC15A (PHY0818A, PhytoAB), and anti-GFP (MBL no. 598). The secondary antibody used was anti-rabbit IgG-HRP conjugate (BIO-RAD). ECL Prime Western blotting detection reagent (Cytiva) was used with ImageQuant LAS 4000 (GE Healthcare) to detect chemiluminescence. Western blotting of all organelle markers and anti-GFP was performed on three independent experiments of flower protein fractionation (except anti-Sec21p, one replicate).

### Nano LC-MS/MS comparative proteomics

Each of the sucrose fraction from 6/12%, 12%, 12/18%, and 18/25% sucrose fraction and interphases were treated with 2% SDS to extract membrane proteins before sending for comparative proteomic analysis service provided by Kazusa DNA Research Institute (Japan).

### Statistical Analyses

Statistical analyses were performed using GraphPad software (Domatics) and Microsoft Excel. For all box plots, center line shows the median, box limits indicate the 25th and 75th percentiles, whiskers extend 1.5 times the interquartile range from the 25th and 75th percentiles and data points are plotted as closed circles. All box plots were generated by BoxPlotR (Spitzer et al., 2014).

### Accession numbers

The Arabidopsis AGI locus identifiers are as follows: *DIC2* (At4g24570), *AtVPS13a* (At1g48090), *AtVPS13b* (At4g17140), *AtSHRUBBY* (At5g24740), and *AtRabA4B* (At4g39990).

## Supplemental data

**Supplemental Figure S1**. Luciferase-based pollen compatibility assay

**Supplemental Figure S2**. Phenotypes of the *atvps13a* mutants isolated in the genetic screen

**Supplemental Figure S3**. Structural conservation of AtVPS13 proteins and their homologs

**Supplemental Figure S4**. Reduced male transmission efficiency of *atvps13a* allele

**Supplemental Figure S5**. DAPI staining of pollen grains from open flowers

**Supplemental Figure S6**. Genotype of genome-edited *atvps13b* in Col-0 background

**Supplemental Figure S7**. RNA-seq counts from a Transcriptome variation analysis database (travadb.org)

**Supplemental Figure S8**. Ectopic vesicle fusion in the hydrated *atvps13a* pollen grains

**Supplemental Figure S9**. Dynamic of relative intensity of AtVPS13a:Venus with mCherry:RabA4B, or R-GECO1 at the polarized site in dual reporter pollen grains

**Supplemental Figure S10**. Putative C2 domain of AtVPS13a is important for efficient pollen germination and its distribution in pollen tube

**Supplemental Table S1**. List of detected proteins in selected sucrose fractions and their Pearson correlation coefficient with AtVPS13a

**Supplemental Table S2**. Result of PANTHER GO-Slim Biological Process analysis of proteins with high (>0.8) Pearson correlation coefficient with AtVPS13a

**Supplemental Table S3**. List of proteins uniquely found in AtVPS13a:Venus Co-IP sample and their Pearson correlation coefficients with AtVPS13a:Venus in sucrose gradient fractions proteomics

**Supplemental Table S4**. List of oligonucleotides used in this study

**Supplemental Table S5**. List of protein databases used to retrieve VPS13 sequence of each species

**Supplemental Movie S1**. Live imaging of *in vitro* pollen germination of dual reporter line AtVPS13a:Venus and mCherry:RabA4B

**Supplemental Movie S2**. *In vivo* pollen Ca^2+^ spike during pollen hydration shown by ratiometric YFP/CFP video of pollen expressing YC3.6 calcium reporter

## Acknowledgements

We thank M. Okamura, T. Manabe, Y. Yamamoto, M. Niidome, M. Ishii, and A. Yoshida for their technical assistance. We also thank M. Takeuchi for providing the cryoporter. This work was supported in part by Grant-in-Aid for Transformative Research Areas (22H05174 to S.F.), Grants-in-Aid for Scientific Research on Innovative Areas (23113002, 16H06467, 16H06464, 21H05030 to S.Tak.; 16H01467, 18H04776 to S.F.), Grants-in-Aid for Scientific Research (25252021, 16H06380 to S.Tak.; 18H02456 to S.F.), Grant-in-Aid for Challenging Exploratory Research (15K14626 to S.F.), Grant-in-Aid for JSPS Fellows (19J01563 to Y.K.; 18J13423 to S.Tan.) from the Ministry of Education, Culture, Sports, Science and Technology of Japan (MEXT). This work was also supported by the PRESTO program (JPMJPR16Q8 to S.F) of the Japan Science and Technology Agency and the Suntory Rising Stars Encouragement Program in Life Sciences (to SF). The authors declare no competing financial interests.

## Author contributions

S.F. conceived the study. S.Tan., S.F., and S.Tak. planned the experiments. S.Tan., S.F., and S.Tak. wrote the manuscript with inputs from all other authors. S.Tan. conducted most of the experiments and data analysis. M.I. helped establish the luciferase assay. F.I. performed the transmission electron microscopy experiments. Y.K. performed the phylogenetic analysis. T.S. conducted the bulk-segregant sequence analysis.

